# The behaviour of hoverfly larvae (Diptera, Syrphidae) lessens the effects of floral subsidy in agricultural landscapes

**DOI:** 10.1101/045286

**Authors:** Elsa A. Laubertie, Steve D. Wratten, Alexandra Magro, Jean-Louis Hemptinne

## Abstract

Modern agricultural landscapes favour crop pests: herbivores benefit from resource concentration and/or the absence of natural enemies in large areas of intensively farmed fields interspersed by small fragments of natural or non-crop habitats.

Conservation biological control (CBC) aims at increasing the functional diversity of agricultural landscapes to make them more hospitable to natural enemies, and less to herbivores. Although natural enemies readily respond to this management, very few studies assess if they succeed in effectively protecting crops.

We set up a field experiment to study if an ecological infrastructure varying in size and consisting of the provision of floral resources at the centre of lettuce plots would influence the number of eggs laid by hoverflies, and ultimately the control of lettuce aphids. We found that the hoverfly females lay more eggs in the plots with the larger flower resource compared to the control. However, this response had no impact on the abundance of aphids on the lettuces.

We designed two laboratory experiments to understand this absence of response. We found mutual interference between hoverfly larvae, and suggest it may undermine the biological control of aphids.

This mismatch between landscape management and the response of hoverflies indicates CBC should take into account insect behaviour, particularly their response to conspecific density, additionally to landscape ecology.

## Introduction

Modern agricultural landscapes display large areas of intensively farmed fields interspersed by small fragments of natural or non-crop habitats (1-3). Crops are susceptible to pest because arable lands are frequently disturbed by cropping practices that impede the development of food webs that may deliver pest regulation (4-6). Herbivorous insects can emigrate from refuges in crop or non-crop habitats and colonize new crops (4, 7), eventually benefiting from the spatial concentration of their host-plants (8, 9). Natural enemies have lower dispersal and reproductive rates; they are disadvantaged if non-crop refuges are rare or absent, and too far from the crops (10). Herbivores may therefore experience an enemy free space (11, 12); that may explain why farmers tend to use more insecticides in simplified landscapes (13, 14) but see (15).

Conservation biological control (CBC) aims at correcting the above mentioned imbalances in favour of herbivores by managing agricultural landscapes to make them more hospitable to natural enemies (5, 16-18). Ecological infrastructures are added to agricultural landscapes to provide natural enemies either with shelters to survive adverse conditions, alternative sources of prey/hosts or nectar and/or pollen (16, 19). Therefore, a stronger numerical response of natural enemies to prey, and a better synchrony between the arrival of prey and natural enemies are anticipated. This should lead to efficient biological control of pests (20).

Natural enemies have a strong positive response to management practices that increase landscape complexity (11, 16, 21). However, very few studies assess if this positive response translates into crop protection (11, 22-25). The few that went as far as measuring the relationships between landscape complexity, natural enemies and crop protection produce contradictory results that are not easily explained (26-28).

Four knowledge gaps are frequently invoked for this absence of pattern: the relative importance of emigration and immigration of natural enemies between crops and non-crop habitats, the lack of information on the birth and death rate of natural enemies in the various compartments of the mosaics of habitats, the effect of crop management practices on the above mentioned processes, and finally the timing, frequency and amplitude of movement between non-crop and crop habitats (3, 10, 22, 27).

To contribute to this debate we firstly set up a field experiment to assess the effects of the abundance of natural enemies while controlling for the confounding effects of all the processes related to emigration and immigration between the various compartments of the agro-landscape. To manipulate the abundance of natural enemies, we provided floral subsidy to aphidophagous hoverflies (Diptera, Syrphidae) by planting buckwheat (*Fagopyrum esculentum* Moench) at the centre of lettuce plots, and we also varied the area of this subsidy. In this manner, all the lettuces were equally closed to buckwheat. We expected that larger subsidy would be positively correlated to the presence of more eggs of hoverflies on the surrounding lettuces infested with aphids. As a consequence, we were expecting fewer aphids on the lettuces planted around the larger subsidy of buckwheat. Secondly, we assessed in the laboratory if the larval density affected the behaviour of hoverfly larvae. We anticipated that the larvae at high density would kill fewer aphids as a result of their interactions with other larvae. Mutual interference reduces predation efficiency of individual natural enemies and is one of the major ecological limits to biological control (29). It occurs when the amount of intraspecific interactions increases with the density of natural enemies up to a point where it reduces the time available for searching and handling prey, eventually triggering aggressive behaviour and cannibalism (29, 30).

We selected this biological model because aphids are among the major pests for several crops of economic importance in temperate regions, including lettuces (31). Hoverflies are natural enemies that have an influence on aphid abundance (32), and they respond to ecological infrastructures such as floral subsidies (33-36).

## Material and methods

### Incidence of added floral resources on hoverfly and aphid abundance in the field

To assess the influence of supplementary pollen and nectar on the abundance of hoverflies and aphids, a field experiment was set up at the Horticultural Research Area (HRA) and at Iverson Field (IF), on the campus of Lincoln University, New Zealand. The HRA comprises 26 ha of land divided into 19 blocks, and IF consists of an area of 13 ha divided into 14 blocks. Five blocks of similar dimensions at the HRA and one at IF were used (Table 1). The 5 blocks at the HRA were located within a circle of 370 m while IF’s was 500 m away to the Southwest. On December 15, 2005, at the centre of each of these six blocks, a 130 × 20 m strip was prepared for sowing. Then, in each of these strips, three 20 × 20 m plots were delineated and separated from each other by 35 m (Fig 1). A mixture of grass species was sown in the areas between the plots. Later, the turf was regularly mown. Those strips were surrounded by herbaceous vegetation briefly described in Table 1. One of the following treatments was randomly allocated to each 20 × 20 m plot: 1) a control only consisting of lettuces (*Lactuca sativa* L. cv. Target) planted at a spacing of 45 cm x 90 cm; 2) lettuces planted as above but the centre of the plot was occupied by a 3 m x 3 m area of buckwheat (*Fagopyrum esculentum* Moench cv. Katowase), and 3) lettuces plus a 12 m x 12 m area of buckwheat at the centre of the plot. Buckwheat provides nectar and pollen, which adult hoverflies readily use (34). It was sown on December 28, 2005 and January 13, 2006; the lettuces were transplanted on January 19 and 20, 2006. This experimental design allows testing if CBC is dependent on the abundance of floral resources while controlling for the confounding issue of natural enemy movement between non-crop and crop habitats.

Buckwheat was indeed sown at the centre of the lettuce plots and all the lettuces were equally distant from the flowers.

**Fig 1.**
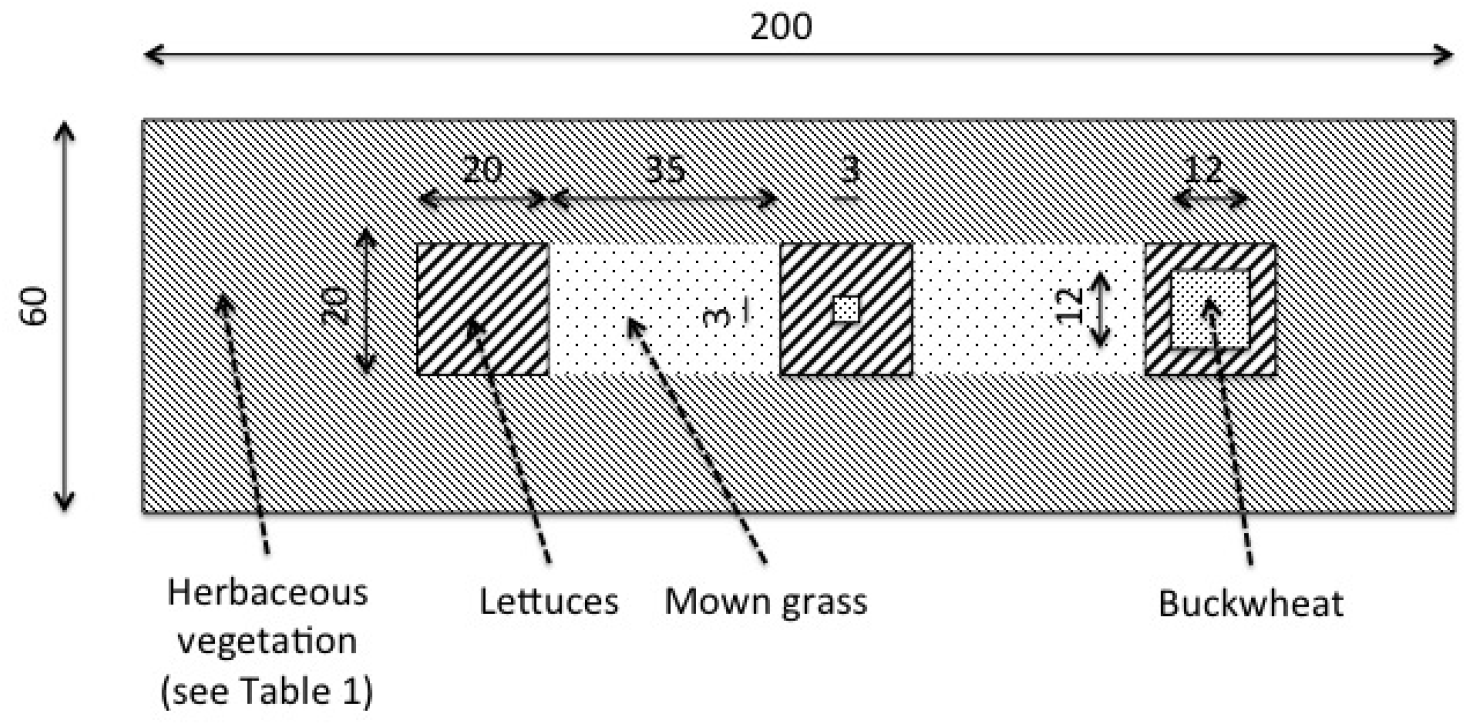
*Plan of one experimental block, showing the dimensions and location of lettuces and buckwheat plots*.

**Table 1.**
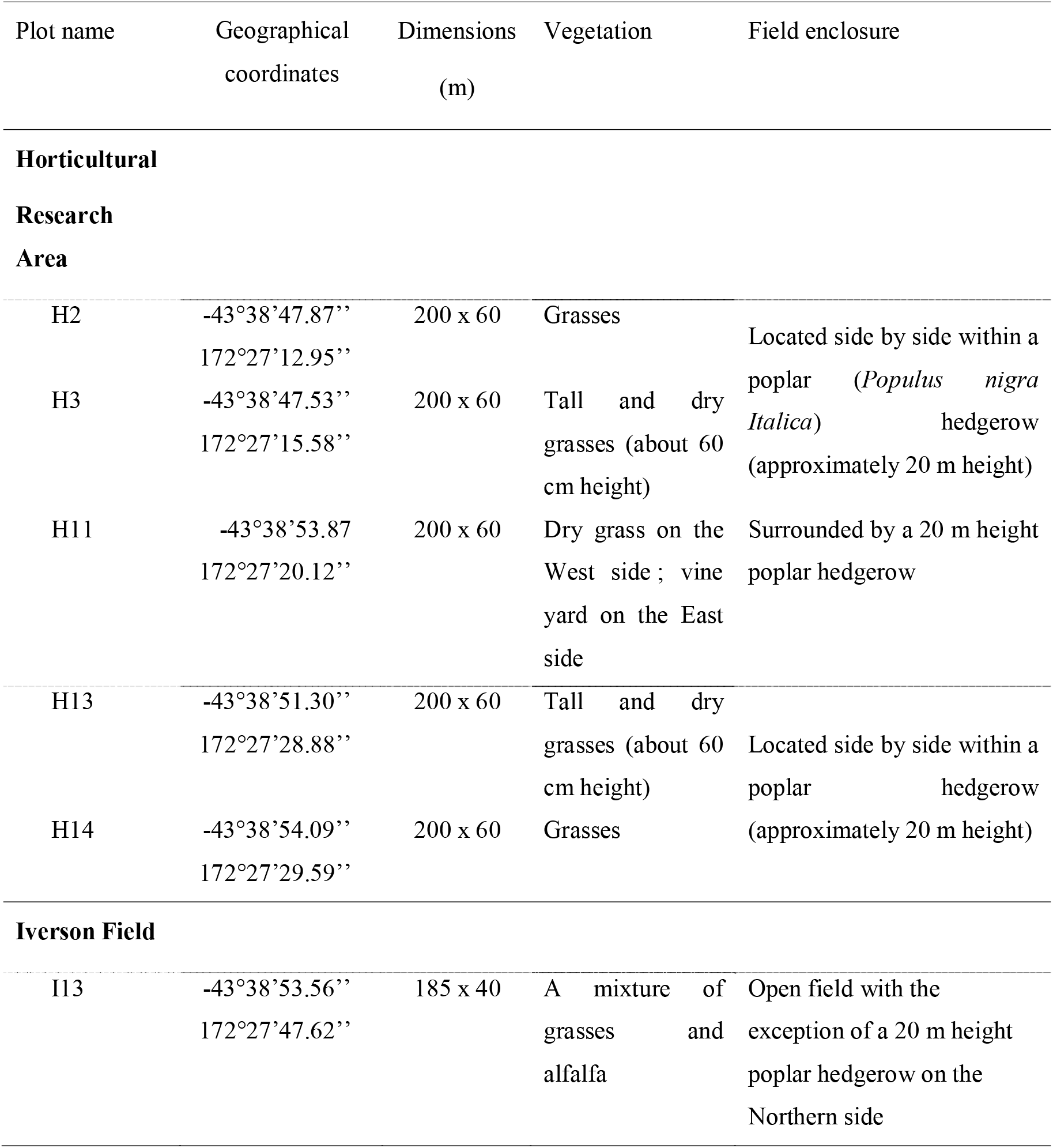
*The geographical coordinates, dimensions, type of vegetation around the experimental area, and information on the field enclosure of the 6 experimental fields*.

Plots were kept free of weeds by hoeing and were visited every day to record the presence of hoverflies. Adult hoverflies were seen for the first time on February 15, 2006, when buckwheat started to flower. Sampling for aphids, hoverfly eggs and larvae started the next day. Six lettuces were randomly selected in each plot and their shoot carefully cut at ground level on February 16, 2006. They were individually put in large plastic bags and brought back to the laboratory. There, each leaf was carefully cut off, unfolded and inspected. The aphids, and the hoverfly eggs and larvae were counted. This was repeated on February 27, March 13, and March 22 when the lettuces reached marketable size.

The numbers of aphids, syrphid eggs and larvae were analysed using General Linear Mixed Models with a Poisson distribution (package lme4 in R 2.11.1). The added floral resource (no buckwheat, small or large area of buckwheat) was a fixed factor. The weeks and blocks were random factors. The weeks were pseudoreplications within each treatment nested in the blocks. *A priori* orthogonal contrasts were implemented to compare the floral subsidies (large and small) to the control, and then the large subsidy to the small (37).

### The behaviour of hoverfly larvae in the laboratory

#### Insect choice and culture

The numerical response of hoverfly females to aphids in the presence of floral subsidy is expected to lead to high density of larvae. Two experiments were set up in the laboratory to see whether mutual interference would appear in such circumstances.

The two dominant species of hoverflies in the experimental fields were *Melangyna novaezelandiae* (Macquart) and *Melanostoma fasciatum* (Macquart). However, we never succeeded in developing a stable and reliable culture of these species in the laboratory to support the experiments. Therefore, we decided to work with *Episyrphus balteatus* De Geer. This choice is firstly justified by the ease of rearing this species, secondly by its phylogenetic relatedness to the two species observed in the field, suggesting ecological resemblance (38-40). *M. novaezelandiae, M. fasciatum* and *E. balteatus* belong to the subfamily Syrphinae ; *E. balteatus* and *M. novaezelandiae* are the most closely related because they are members of the tribe Syrphini while *M. fasciatum* belongs to the Bacchini tribe (41). Because phylogenetic conservatism of interactions occurs in many taxa (42), we believe that knowledge on the behaviour of *E. balteatus* in the laboratory will prove useful to predict how *M. novaezelandiae* and *M. fasciatum* behave in similar conditions.

Adult *E. balteatus* were reared in mesh cages (40 × 75 × 50 cm) in a greenhouse, from April to September 2006. Hoverflies were offered every other day fresh flowers from the field in water. Broad bean plants (*Vicia faba* L.) infested with pea aphids (*Acyrthosiphon pisum* Harris) were periodically introduced into the cages to induce the hoverflies to lay eggs. The beans were checked every other day and the leaves with eggs were removed and kept in the laboratory at 21 ± 1ºC, under a photoperiod of LD 16:8 h. Larvae were reared in the same environmental conditions, in 175 cm³ plastic boxes that contained a piece of corrugated filter paper to reduce the frequency of encounter and therefore the risks of cannibalism. Three times a week the larvae were fed an excess of mixed instars of *A. pisum*. Two cut stems of broad bean were added to each box to improve the survival of the aphids. As hoverfly larvae are active at night, eggs were incubated and larvae reared under a reversed photoperiod to allow for observations during normal working hours. They were kept in darkness from 8:00 to 16:00 and all the observations were made under a red light.

#### Experiment 1. The effects of aphids and the density of hoverfly larvae on larval dispersal

If mutual interference happens, we expect larvae to become more active and to disperse when their density relative to prey availability increases. To assess the tendency of larvae to disperse in the presence of conspecifics and different numbers of prey, a third instar larva (the “focal larva”) of *E. balteatus*, which had been starved for 2 h prior to the experiment, was placed at the centre of a 15 cm diameter Petri dish on a 3 cm piece of broad bean stem. Then, second instar larvae were gently put on the piece of broad bean stem in four treatments described below and repeated 20 times: 1) two conspecific second instar larvae, 2) two second instar larvae and 40 pea aphids, 3) eight second instar larvae, and 4) eight second instar larvae and 40 pea aphids. Larvae of the second instar were used to distinguish the focal larva from those just there to manipulate the density. Every 30 minutes for 2 hours, we recorded if the focal larvae were on the broad bean stem or not. The experiment was performed between 8:00 and 16:00 at 21°C, and under a photoperiod of LD 16:8 h.

The data were binary and there was temporal pseudoreplication. We used two Generalised Linear Mixed Models with binomial errors (package lme4 in R 2.11.1) the first had an interaction between the two independent variables (numbers of larvae, and presence/absence of aphids) while there was no interaction in the second. We tested significance by deletion of the interaction, and compared the change in deviance with a χ^2^ test (Crawley, 2007).

#### Experiment 2. *Mutual interference between* E. balteatus *larvae*

A 3 cm piece of broad bean stem was placed at the centre of a 15 cm diameter Petri dish, along with 2, 8, 16 or 32 second instar larvae of *E. balteatus*, and 40 similar sized pea aphids. Then, a third instar larva of *E. balteatus* (the “focal larva”) that had been starved for 2h prior to the experiment was gently put on the stem. The Petri dishes were continuously and sequentially observed for 30 min and the number of aphids the focal larvae attacked was recorded. Each treatment was repeated 10 times at 21°C, and under a photoperiod of LD 16:8 h, between 8:00 and 16:00. The same observer carried out three replicates of each treatment simultaneously. The searching efficiency of third instar larvae (the number of aphids captured/larva in 30 min) was calculated. The regression of the log of the searching efficiency on the log of the larval density was calculated. The slope *m* is expected to vary from 0 to −1 indicating a growing mutual interference (30).

## Results

### Incidence of added floral resources on hoverfly and aphid abundance in the field

We recorded 51,745 lettuce aphids *Nasonovia ribisnigri* (Mosley) and 284 black bean aphids *Aphis fabae* (Scopoli) on the lettuces throughout the sampling period. We counted 9,257 and 798 hoverfly eggs and larvae respectively. We were not able to assign them to the species but we only spotted adults of *M. novaezelandiae* and *M. fasciatum*.

The mean number of aphids per lettuce was higher at the beginning of the sampling campaign and declined thereafter (Fig 2). Throughout the sampling period the lettuces hosted similar numbers of aphids whether they grew in a plot without or with buckwheat (Contrast_Control versus Treatment plots_: z value=1.240; P=0.215; Contrast_Large versus small plots_: z value=0.510; P=0.610). They were equally infested on the last date when they reached marketable size (Contrast_Control versus Treatment plots_: z value=0.459; P=0.646; Contrast_Large versus small plots_: z value=-0.737; P=0.461).

Some hoverfly eggs were already recorded on February 16. Over the entire campaign, the number of eggs on the lettuces of the treatment plots tended to be more abundant than in the control. However, the difference was not significant (Contrast_Control versus Treatment plots_: z value=-1.860; P=0.063; Fig 3a). The lettuces from the plots with the larger area of buckwheat at the centre had significantly more eggs than those from the plots with the small area of buckwheat (Contrast_Large versus small plots_: z value=2.803; P=0.005; Fig 3a).

**Fig 2.**
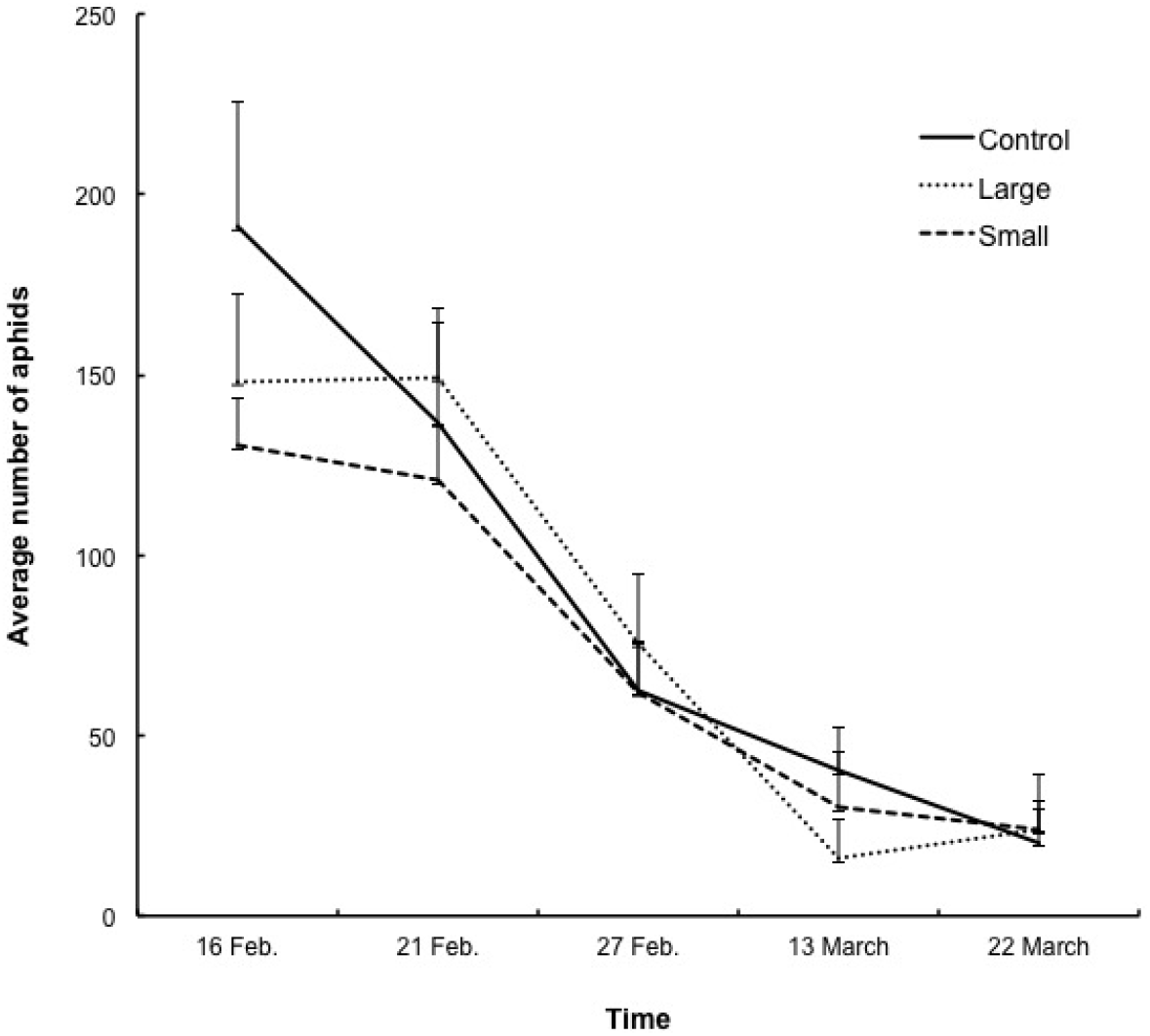
*The average number (SD) of aphids on the lettuces of the experimental plots with a small (3x3 m) or large (12x12 m) area of buckwheat at the centre, or without buckwheat (control) at five successive dates. For clarity, one-sided SD are represented*.

Hoverfly larvae peaked in abundance on February 27, 2006. The number of larvae per lettuce was on average an order of magnitude smaller than the number of eggs. The average number of hoverfly larvae per lettuce in the control plots was not significantly different from the numbers observed in the treatment plots (Fig 3b). However, the large plots had significantly more larvae than the small plots (Contrast_Control versus Treatment plots_: z value=-1.416; P=0.157; Contrast_Large versus small plots_: z value=2.170; P=0.030).

**Fig 3.**
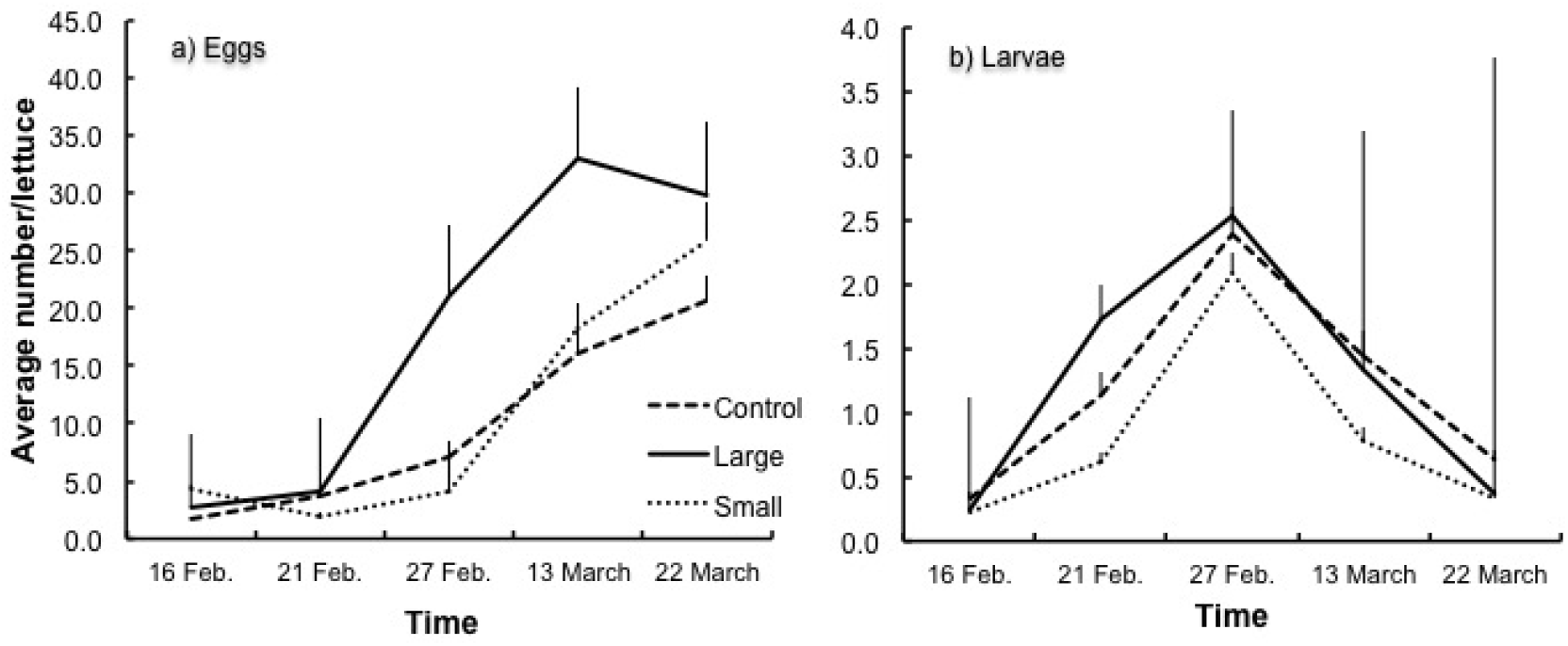
*The average number (SD) of hoverfly eggs (a), and larvae (b) per lettuce in the experimental plots with a small (3x3 m) or large (12x12 m) area of buckwheat at the centre, or without buckwheat (control) at five successive dates. For clarity, one-sided SD are represented*.

### The behaviour of hoverfly larvae in the laboratory

#### Experiment 1. The effects of aphids and the density of hoverfly larvae on larval dispersal

In the first 30 min of the experiment in which the third instar larva interacted with 2 second-instar larvae and 40 pea aphids, the proportion of third instar larvae on the broad bean dropped from 80 to 50%, and then remained at that level for the rest of the experiment. On the contrary, in the three other treatments, the proportions were much lower in the beginning: 40% and even 25% in the treatment with 8 second-instar larvae and no aphids. Then, it steadily declined over the course of the experiment. After 2 h, only 5 to 15% of the third instar larvae, depending on the treatment, still were on the broad bean (Fig 4).

**Fig 4.**
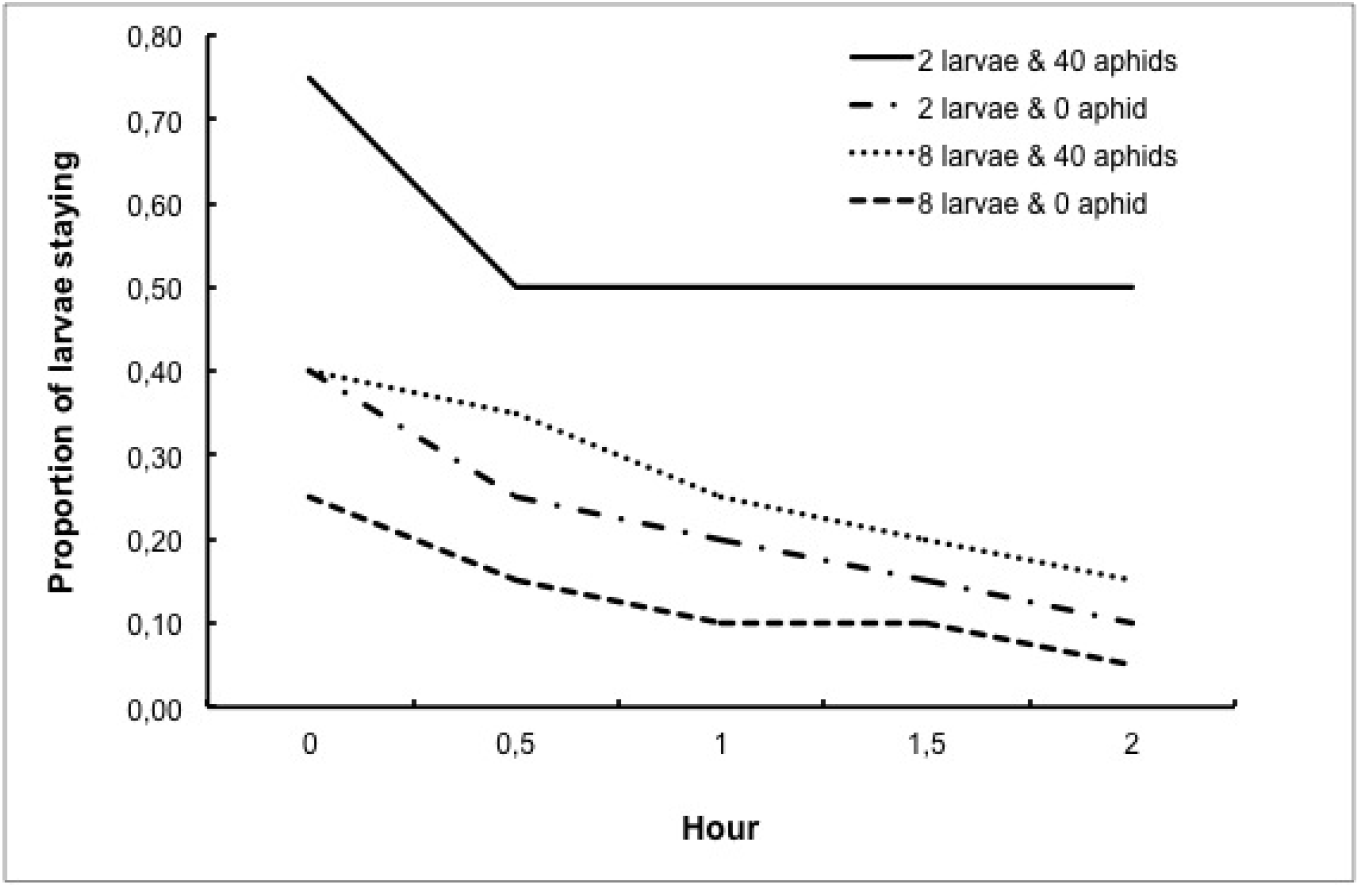
*The proportions of third-instar larvae of Episyrphus balteatus staying on a piece of broad bean stem in the presence of 2 second-instar conspecific larvae, 2 second-instar conspecific larvae and 40 pea aphids, 8 second-instar conspecific larvae or 8 second-instar conspecific larvae and 40 pea aphids throughout an observation period of 2 hours*.

The interaction between the presence/absence of aphids and the density of second instar larvae was not significant (z value=-1.674; P=0.094). The presence of aphids, contrary to the number of 2^nd^ instar larvae, had a significant effect on the number of 3^rd^ instar larvae staying on the broad bean stem (Presence/absence of aphids: z value=2.972; P=0.003; N^r^ of 2^nd^ instar larvae: z value=-0.426; P=0.671).

#### Experiment 2. *Mutual interference between* E. balteatus *larvae*

The value of the attack rate of the third-instar larvae gradually declined when the density of second instar larvae increased (Table 2).

**Table 2.** *The mean number of aphids eaten, and the attack rate (mean number of aphids eaten.larva^-1^.30 min^-1^) by third-instar larva of Episyrphus balteatus in the presence of 2, 8, 16 or 32 second-instar conspecific larvae*

| Nr of<br>2<sup>nd</sup>-instar larvae | Mean nr eaten<br>(min-max) | Attack rate |
| --- | --- | --- |
| 2 | 7.6 (6-9) | 2.53 |
| 8 | 21.7 (15-29) | 2.41 |
| 16 | 29.3 (19-33) | 1.72 |
| 32 | 35.7 (33-37) | 1.08 |

The slope of the relationship between the log of the searching efficiency (the number of aphids captured/larva/30 min) and the log of the predator density (Fig 5) is significantly different from zero (y = −0.4269 x – 0.1903; F = 47.01; df = 38; P < 0.05), which indicates the existence of mutual interference between larvae.

**Fig 5.**
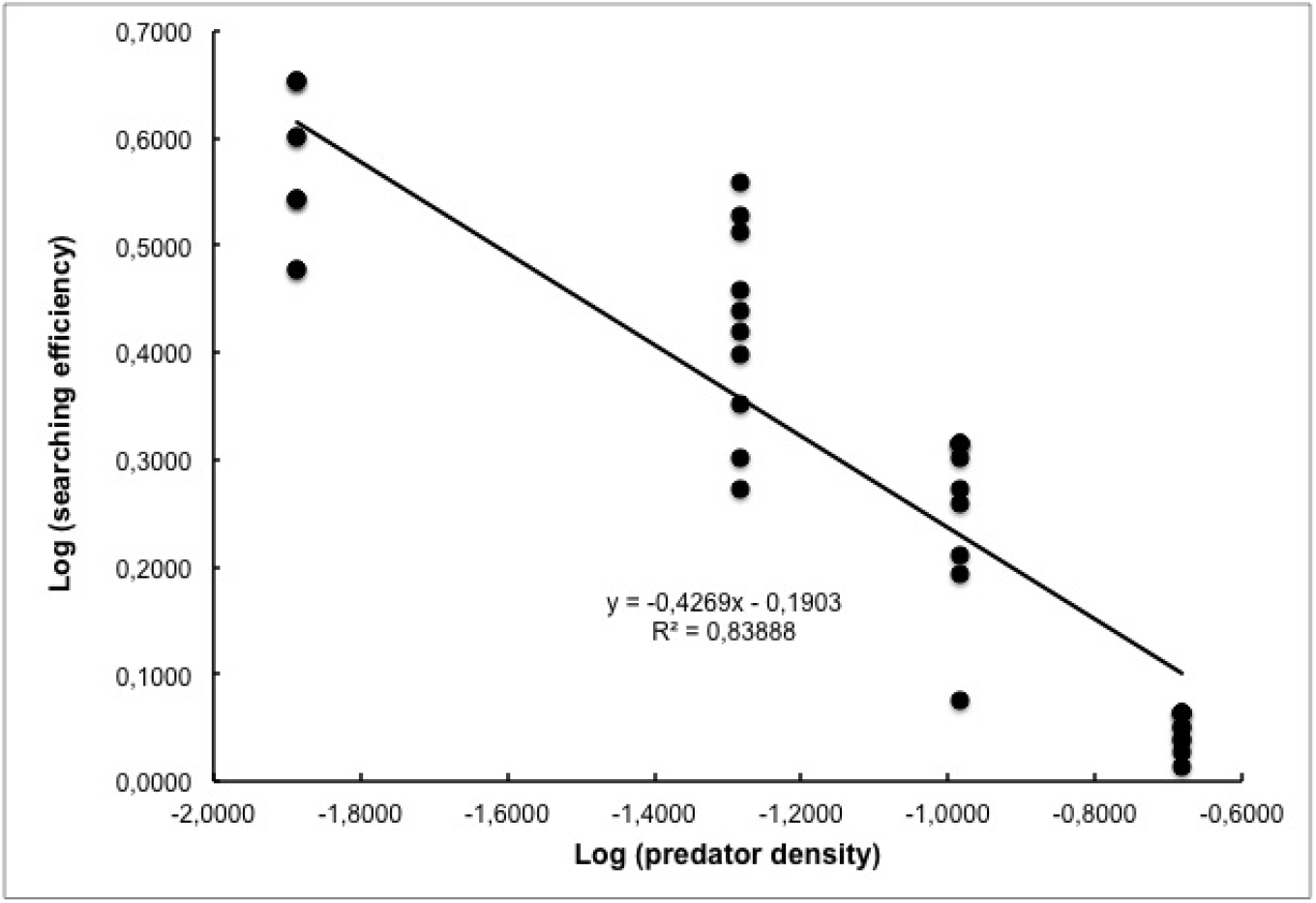
*The searching efficiency (aphids captured.unit time^-1^.hoverfly larva^-1^) of third-instar larvae of Episyrphus balteatus as a function of the density of second-instar conspecific larvae*

## Discussion

In our field experiment, we observed more eggs of hoverflies on the lettuces planted around large floral patches of buckwheat than on the lettuces in the control. There was also a gradient in the egg numbers from the lowest in the control plots, intermediate in the plots with a small area of buckwheat to the highest in the plots with the largest area of flowers. Buckwheat is known to attract several species of hoverflies to which it provides nectar and pollen in quantity and quality (34, 43). Our results support the practice of planting sweet alyssum (*Lobularia maritima* (L.) Desv.) in organic lettuce fields to attract hoverflies (44-47). Our results also confirm that natural enemies respond positively to the provision of additional food sources in crop and non-crop habitats (21).

However, we observe that these higher numbers of eggs did not translate in higher numbers of larvae throughout the experiment or at the end, when the lettuces were harvestable. An explanation would be that hoverflies might be limited in the field by predation or cannibalism in a density-dependent manner (48, 49). Alternatively, mutual interference can be at the origin of the levelling out in the number of larvae (29).

We set up laboratory experiments to see whether hoverfly larvae develop mutual interference when their density becomes too important. We observed that the larvae of *E. balteatus* were highly mobile when aphids were rare. They displayed a strong mutual interference that reduces their searching efficiency for prey. Therefore, if our observations with *E. balteatus* are representative of the behaviour of the two dominant species recorded in our field experiment, as would suggest the theory on niche conservatism (Gomez et al. 2010), the collapse in the numbers of larvae that we observed in the field is likely to be due to the mutual interference between hoverfly larvae.

Finally, aphids were not less abundant on the lettuces around large areas of buckwheat than on those from the plots with only a small area of it or without buckwheat. The changes in the numbers of aphids throughout the field experiment were similar in the three treatments. Also, when the lettuces reached marketable size, the number of aphids per lettuces did not differ significantly across the three treatments, and was still much higher than the economical threshold of damages (50). Contrary to a field cage experiment (Hogg et al., 2011) and a field experiment manipulating the number of hoverfly eggs on lettuces (51), our field experiment shows that an increased abundance of natural enemies in response to the provision of ecological infrastructures failed to reduce aphid abundance.

We did not design our field experiment with an applied perspective in mind. By placing squares of buckwheat at the centre of plots of lettuces we wanted to disentangle the influence of the abundance of floral subsidy from the many factors affecting the movement of those enemies from the locations of the subsidies to the crops throughout the landscape matrix.

In conclusion, actions undertaken to attract natural enemies nearby crops do not always succeed in reducing pest abundance under economic threshold of damage. According to knowledge and expertise accumulated so far, these actions deliver the expected results in terms of predator community composition and population abundance (Landis et al. 2000; Bianchi et al. 2006; Chaplin-Kramer et al. 2011). However, they do not translate in a predictive way in the biological control of pests because they are curtailed by evolutionary trade-offs that shape the life history of plants and insects.

Recent reviews suggest that predictive and efficient conservation biological control is still out of reach for two main kinds of reasons: firstly, the lack of understanding in the movements of natural enemies between the various habitats of agricultural landscape, and secondly the difficulty in estimating demographic rates of natural enemies in relation to habitat types (3, 10, 27, 52). However, these knowledge gaps do not explicitly refer to the behavioural decisions made by individual natural enemies searching for preys or hosts. Without clear insight on the decisions modulating feeding and non-feeding interactions between natural enemies and their preys/hosts the relationship between the sum of individual behaviour and population dynamics will remain out of reach (53). From a pest control point of view, it is probably more important to know how species interact rather than the number and diversity of species in communities (6). We believe that the absence of a food-web perspective coupled to behavioural ecology is probably a largely underestimated knowledge gap that affects the ability to predict the relationship between landscape management and biological control (6, 54). The mismatch between landscape management and the impact of this management for crop level protection does not mean that biological control is not feasible. It simply indicates it still is a complex issue (55).

## Acknowledgments

Shona, J. Randell, S. Clearwater, S. Blyth, D. Riding helped with sampling insects in the field; J-F. Garrigues managed the greenhouse. A. Lister checked the statistical analyses. A. F. G. Dixon made suggestion to improve the manuscript and edited it. Financial support came from the New Zealand Foundation for Research, Science and Technology (LINX 0303). EL and JLH benefited from a Dumont d’Urville grant (The Royal Society of New Zealand and The French Embassy in New Zealand). The Labex TULIP (ANR −10-LABX-41) funded JLH and AM.

